# Siglecs in the Porcine Oviduct and Sialylated Ligands on Sperm: Roles in the Formation of the Sperm Reservoir

**DOI:** 10.1101/2023.03.26.534240

**Authors:** Leonardo M. Molina, Lauren E. Pepi, Asif Shajahan, Kankanit Doungkamchan, Parastoo Azadi, Daniel B. McKim, David J. Miller

## Abstract

During mammalian insemination, most of the deposited sperm are lost by retrograde flow or the female reproductive tract’s immune response. Once semen enters the uterus, seminal fluid and sperm elicit leukocyte infiltration that contributes to the elimination of sperm in the uterus. However, unlike the uterus, invading sperm do not trigger a phagocytic response in the oviduct in the absence of dysfunction or disease states. Thus, the oviduct possesses a distinct immunological microenvironment that tolerates sperm while maintaining the capacity to respond to pathogens. It has been suggested that sperm glycocalyx contributes to innate oviductal tolerance, but the cell and molecular mechanisms are not understood. The current investigation focused on the role of sialic acid-containing glycoconjugates on sperm and their potential to elicit innate tolerance via cognate sialic acid-binding immunoglobulin-type lectins (Siglecs) expressed in the oviduct. In this manuscript, we report our discovery of eight Siglecs (Siglecs-1, -2, -3, -5, -10, -11, -14, -15) expressed in the lower pig oviduct, five of which are known for immune inhibitory functions (Siglecs-2, -3, -5, -10, and -11) and how these may play a role in achieving sperm-induced immune suppression in the oviduct microenvironment. Mass spectrometry profiling of porcine sperm revealed the presence of a mixture of α2,3 and α2,6 linked sialic acids with α2,3-linked sialic acids as the dominant linkage. Of the detected glycans, several sialic acid-containing glycoconjugates were identified as potential ligands for Siglecs (among O-linked glycans: NeuAc_1_GalNAc_1_, NeuGc_1_GalNAc_1_, NeuAc_2_Gal_1_GalNAc_1_; attached to glycolipids: NeuAc_2_Gal_1_GalNAc_1_Gal_1_Glc_1_, Fuc_1_Gal_1_GalNAc_1_NeuAc_1_Gal_1_Glc_1_). This is the first report of Siglec expression in the mammalian oviduct and total glycan analysis of porcine sperm. The results of this study reveal the potential for a sperm-sialoglycan and oviductal-Siglec axis that may contribute to the distinct immunophysiology of the oviduct fundamentally required for undisrupted reproduction in mammals.

## Introduction

After semen deposition in mammals, only a small subpopulation of the deposited sperm can successfully ascend beyond the uterus and reach the site of fertilization in the upper oviduct (Suarez & Pacey, 2006). A large portion of the deposited sperm in swine and other species is often lost due to uterine backflow (Hernández-Caravaca et al., 2012). Further sperm clearance is mediated by the female reproductive tract’s local innate immune response. Interactions between the sperm, seminal plasma, and/or semen extenders with the uterine milieu mediate an immediate inflammatory response which results in the release of pro-inflammatory cytokines and chemokines such as IL-1, IFN-γ, TNF-α, and IL-8 (Katila, 2012).

The release of these inflammatory mediators in the uterus stimulates the recruitment of leukocytes, mainly polymorph neutrophils (PMNs), to the uterine lumen (Katila, 2012). The recruited leukocytes aid in the rapid clearance of sperm from the uterus, theoretically preventing the production of sperm-specific antibodies (Katila, 2012). There is evidence that certain females express antibodies against sperm antigens resulting in lower fertility rates. This highlights an additional mechanism that could play a role in sperm clearance from the female reproductive tract in some females (Ghaderi et al., 2011).

Sperm that reach the oviduct can adhere to the lower oviduct region, known as the isthmus, to form the oviductal sperm reservoir. Sperm binding to the oviduct suppresses the elevation of intracellular Ca^2+^ and slows sperm motility (Dobrinski et al., 1996, 1997). Subsequently, sperm are gradually released from the reservoir by several possible mechanisms (Sharif et al., 2021, 2022) and move toward the site of fertilization in the ampulla. The gradual release of sperm is thought to help control polyspermy rates (Hunter & Léglise, 1971). The ability to prolong sperm lifespan lengthens the fertilization window, which is particularly important in species in which mating and ovulation are not tightly synchronized (Birkhead & Møller, 1993; Miller, 2018). The formation of the sperm reservoir is a result of sperm binding to the oviduct epithelium in a carbohydrate-mediated fashion. Glycans found on oviduct epithelial cells act as ligands for sperm surface receptor proteins (Suarez, 2001). Previous studies from our lab have shown that porcine sperm preferentially bind to two oviduct glycan motifs, a 6-sialylated multi-antennary oligosaccharide and 3-O-sulfated Lewis X trisaccharide (Kadirvel et al., 2012; Machado et al., 2014). The binding of sperm to these immobilized glycans in vitro can prolong sperm viability by regulating intracellular Ca^2+^ (Machado et al., 2020), reducing ubiquinone precursor, diminishing electron transport chain activity, and reducing production of reactive oxygen species (Hughes et al., 2023).

Oviduct secretions and signaling provide an optimal microenvironment to support the prolonged survival of sperm throughout the duration of the sperm reservoir. The porcine oviduct is known to contain few PMNs and macrophages regardless of the stage of estrus (Jiwakanon et al., 2005). This is especially true of the isthmus as compared to other regions of the oviduct (Jiwakanon et al., 2005). Furthermore, unlike the uterus, sperm infiltration into the oviduct does not result in the recruitment of phagocytic immune cells (Jiwakanon et al., 2006). The oviduct can tolerate allogenic sperm and the semi-allogenic conceptus as they transit and/or are stored in the oviduct while still being able to respond to dysfunction or disease states. In fact, the opposite appears to take place as sperm-oviduct interactions have been shown to induce an immunosuppressive response. The incubation of human sperm with oviduct epithelial cells in vitro results in the downregulation of the gene expression of pro-inflammatory interleukins such as IL-16, IL-17, IL-8, IL-1A, and IL-1B and pro-inflammatory chemokines including CCL-3, CCL-8, CCL-11, CCL-20, CCL-24, CXCL-9, and CXCL-13 (Mousavi et al., 2021). These and other immunosuppressive signals stimulated by sperm interactions with oviduct epithelial cells could prevent the local recruitment of immune cells and thus prevent sperm from being targeted by the immune system.

Cell surface glycans are known to be major modulators of the innate immune response. Sialic acids are nine-carbon sugars and are often found on the termini of cell surface glycans in mammals (Schauer, 1982). Sperm from multiple species have been shown to possess a thick glycocalyx with a high abundance of sialoglycans (Pang et al., 2007; Wang et al., 2018). It is estimated that approximately 40% of the monosaccharides on the surface of human sperm are sialic acids (Calzada et al., 1994). The composition of the sperm sialome is dynamic and events such as spermatogenesis, transit through the epididymis, incubation with accessory gland fluids, and sperm capacitation are known to remodel the sperm sialome (Ma et al., 2012; Ma et al., 2016). The establishment of proper sperm sialylation is critical for sperm survival and functions. For instance, changes in the sperm sialome can alter the degree of sperm phagocytosis by uterine macrophages in vitro (Ma et al., 2016). Mouse epididymal sperm that had their sialic acids enzymatically removed experienced higher rates of phagocytosis by uterine macrophages than untreated sperm (Ma et al., 2016). Surprisingly, epididymal sperm that gained additional sialic acids by exposure to accessory gland fluids also underwent higher rates of phagocytosis when compared to the control epididymal sperm (Ma et al., 2016). This suggests that sperm require an appropriate intermediate abundance or linkage of sialic acid residues and that the female reproductive tract possesses a distinct immunological microenvironment capable of distinguishing the type and quantity of sialoglycans present on sperm to adequately mount an immune response.

Sialic acid-terminating glycans interact with several proteins, the most abundant of which are Siglecs. Siglecs are proteins involved in self-recognition and combatting pathogens. Siglecs are broadly expressed on cells from the innate and adaptive immune systems and can regulate immune cellular immune activation signaling pathways. Additional roles include the mediation of cell-to-cell adhesion, the attraction of immune cells, and the facilitation of pathogen internalization (Chang & Nizet, 2014; Varki & Angata, 2006). Siglecs have also been characterized in non-immune cells and tissues. For instance, Siglec-12 has been found on epithelial cells of the kidney tubules and prostate epithelium (Mitra et al., 2011). Moreover, recent studies have discovered certain Siglecs on the human and mouse endometrium (Tecle et al., 2019) and cervical epithelium (Landig et al., 2019). It has been proposed by Tecle and colleagues that sperm interact with Siglecs as they transverse the uterus and that Siglec-induced signaling could help sperm survive the severe inflammatory response that takes place in the endometrium following insemination (Tecle et al., 2019). However, there is no conclusive evidence of Siglec function in sperm immune evasion or sperm-induced immune suppression through Siglec signaling. Nevertheless, a study on *Neisseria gonorrhea* infections has shown a positive correlation between the availability of immune activating Siglecs in the cervical epithelium and the ability of patients to combat gonorrhea infections (Landig et al., 2019). Furthermore, it has been shown that mouse sperm can bind to certain recombinant Siglec proteins in vitro (Ma et al., 2016). Together, this shows the potential for Siglecs to mediate immune signaling in response to sialylated pathogens and allogenic cells inside the female reproductive tract. The biasing of the oviduct microenvironment towards immune inhibition in response to sperm interactions is still not well understood. We hypothesize that the interactions between sperm and the oviduct epithelium allow for the formation of the sperm reservoir through sialoglycan-Siglec-mediated interaction and their resulting downstream signaling.

## Materials and Methods

### End-Point RT-PCR

The curated mRNA sequence for Siglec-1 and the predicted sequences for Siglecs-2, -3, -4, -5, -10, -11, -13, -14, and -15 were recovered from NCBI’s GenBank database. We did not look for Siglec-4 as it is not found in hematopoietic cells and is exclusive to nervous tissues (Lehmann et al., 2004). Oligonucleotide primers for each gene were designed using NCBI’s Primer-BLAST online software and manufactured by Invitrogen. All primer sequences used in this study are listed in **Supplemental Table 1**. Whole female reproductive tracts from both pre- and peri-pubertal gilts were collected from Rantoul Foods (Rantoul, Illinois), and sow reproductive tracts were collected from Calihan Pork Processors Inc (Peoria, Illinois). No experiments on live animals were performed in this study. The cyclic stage of the animals was determined based on ovarian morphology (Carrasco et al., 2008). The isthmus region of the oviduct was excised and placed in PBS on ice. Primary oviduct epithelial cells (OECs) were collected by scrapping a glass slide across the outside of the isthmus. OECs were then centrifuged at 5000 x g for 1 min and resuspended in RNAlater (Applied Biosystems) and stored at -20°C until RNA extraction. Total RNA was extracted from OECs and spleen tissues using the RNeasy Mini Kit (Qiagen) following the manufacturer’s guidelines. The optional on-column DNase digestion step was performed to remove any genomic DNA contamination from samples. RNA integrity was assessed by loading 100 ng of total RNA in a 1% agarose gel (Wt/Vol) resolving for 70 min at 70 volts and assessing the 28S and 18S ribosomal bands. RNA purity and concentrations were then assessed based on the absorbance readings measured using a NanoDrop 1000 Spectrophotometer (Thermo Fischer Scientific). Only samples with a 260 nm/280 nm absorbance ratio ≥ 1.9 were used for PCR reactions. RT-PCR reactions were performed using the OneTaq One-Step RT-PCR Kit (New England BioLabs) following the manufacturer’s instructions. A final reaction volume of 25 µL with 100 ng of total RNA and a final concentration of 200 nM oligonucleotide primers were utilized in each reaction. RT-PCR reactions were performed in a SimpliAmp Thermal Cycler (Applied Biosystems) following the cycling conditions described in **Supplemental Table 1**. To evaluate the size of PCR products, 6 µL of each sample were resolved by gel electrophoresis in a 2% agarose gel in TAE buffer (Wt/Vol) for 90 min at 90 volts. A Quick-Load 1kb DNA Ladder or a Quick-Load Low Molecular Weight Ladder (New England BioLabs) was used to estimate the PCR product sizes and confirm the presence of the target genes. The housekeeping gene *GapdH* was used as a loading control for all samples. No template controls (NTC) were also performed for each target gene and no amplification product was observed. All experiments were performed in triplicate in different biological samples. Ethidium bromide signals were imaged in an ImageQuant LAS 4000 (GE Healthcare Biosciences). The raw image was then inverted using the ImageQuant TL software (GE Healthcare Biosciences) and saved as a bitmap file.

### Processing of Sperm

Porcine semen was provided by PIC (Hendersonville, TN) and was stored at 17 °C for up to 24hr before use. Sperm motility parameters were assessed using the Hamilton Thorne Semen Analysis CASA system (Hamilton Thorne). Only samples with initial total motility of >80% were used for experiments. Two mL of semen were washed at 600 x g for 8 min in 10 mL of dmTAGP (2.0 mM CaCl_2_, 4.8 mM KCl, 1.2 mM MgCl_2_, 95 mM NaCl, 1.2 mM KH_2_PO_4_, 25 mM NaHCO_3_, 1 mM pyruvic acid, 5.56 mM glucose, 15 mM HEPES, 0.6% BSA, pH 7.3 – 7.4, sterile filtered). Sperm were resuspended in 5 mL of dmTAGP and washed again at 600 x g for 5 min. The sperm pellet was re-suspended in 1 mL of dmTAGP and the sperm concentration was determined using a Neubauer hemocytometer.

### Protein Extraction

Five hundred micrograms of sperm were washed with PBS at 3000 x g for 5 min and the supernatant was discarded. The samples were dissolved in urea lysis buffer (8 M urea, 200 mM Tris HCl, 100 mM DTT) and incubated at 45°C for 60 min. Following incubation, the samples were centrifuged at 14000 x g for 15 min and the supernatant was passed through a 10 kDa molecular weight cut-off (MWCO) Amicon Ultra centrifuge filter (Millipore Sigma). Pellets were washed with 50 mM ammonium bicarbonate and pelleted once more. The supernatant was added to the MWCO filter. This process was repeated once more. The sample retained by the filter was removed and combined with the cell pellet. The extracted protein samples were homogenized with 1 mL of 50 mM ammonium bicarbonate.

### Glycosyl Composition and Fatty Acid Analysis

Glycosyl composition analysis was performed by combined gas chromatography/mass spectrometry (GC-MS) of the per-O-trimethylsilyl (TMS) derivatives of the monosaccharide methyl glycosides produced from the sample by acidic methanolysis as described previously by Santander et al. (Santander et al., 2013). Briefly, the extracted protein pellet samples were heated with methanolic HCl in a sealed screw-top glass test tube for 18 h at 80°C. After cooling and removal of the solvent under a stream of nitrogen, the samples were derivatized with Tri-Sil (Pierce) at 80 °C for 30 min. GC/MS analysis of the TMS methyl glycosides was performed on an Agilent 7890A GC interfaced to a 5975C MSD, using a Supelco Equity-1 fused silica capillary column (30 m X 0.25 mm ID).

### N- and O-Glycan Release

Approximately 100 µg of extracted protein was incubated with 2 µL (1,000 U) of PNGase F (New England Biolabs) at 37°C for 48 hrs. An additional 2 µL of PNGase F was added after the initial 24 hrs. Following PNGase F digestion, the samples were pelleted, and the supernatant was passed through a 10 kDa MWCO filter. The pellet was washed with ammonium bicarbonate and the supernatant was passed through the 10 kDa filter. This was repeated once more. The released N-glycans (filtrate) were pooled and loaded onto a C18 cartridge (Resprep, 26030) and eluted with 5% acetic acid and lyophilized. Lyophilized samples were then permethylated as described below. The sample remaining in the filter, which contains the O-glycoprotein portion, was combined with the cell pellet. The O-glycoprotein portion was lyophilized and then subjected to reductive β-elimination. Two hundred and fifty µL of 50 mM sodium hydroxide (NaOH) solution was added to the dried sample. The pH was checked to confirm alkaline conditions. Nineteen milligrams per 250 µL sodium borohydride (NaBH_4_) in 50 mM NaOH was added, the sample was vortexed, and then incubated at 45°C for 18 hrs. The sample was then removed from heat and allowed to cool to room temperature. The sample was neutralized by adding 10% acetic acid dropwise until formation of bubbles stopped. The neutralized sample was then loaded onto a packed ion exchange resin (DOWEX H+) in a poly-prep chromatography column (Bio-Rad, 731-1550). The column was then washed with 5% acetic acid. The flow-through was then loaded onto a C18 column and eluted with 5% acetic acid. The sample was then lyophilized. Borates were removed by adding 500 µL of MeOH: Acetic acid (9:1) and drying under a stream of nitrogen. This process was repeated 5 times. Dried samples were then permethylated as described below.

### Glycolipid extraction and purification

An additional aliquot of the purified protein was used for glycolipid extraction. Lipids were extracted using a Folch method (C/M/W= 4/8/3 v/v/v). The supernatant was removed, and the process was repeated 2 times. The supernatant was combined and dried under a stream of nitrogen. Phosphoglycerolipids were removed by saponification. One milliliter of 0.5 M KOH in MeOH (MeOH:water 95:5 v/v) was added to the lipid sample and incubated at 37°C overnight. Following incubation 1 mL of 5% acetic acid was added to the reaction on ice. The samples were then desalted with a tC18 sep-pak cartridge (Waters Corp) and eluted with MeOH. The sample was dried under a stream of N2. Free fatty acid removal was then done by adding 200 µL of hexane to the sample, vortexed, and spun down. Following pelleting the hexane was removed, and this process was repeated 2 times. Lipid pellets were then dried down and permethylated as described below.

### Per-O-Methylation

A dimethylsulfoxide (DMSO)/NaOH base was made by combining 100 µL 50% NaOH (v/v) and 200 µL MeOH in a glass tube and vortexing. Four mL of anhydrous DMSO were then added to the solution and mixed vigorously. The solution was then centrifuged at 3000 x g for 5 mins. The white solid precipitate that formed at the top of the solution and all remaining DMSO was removed. This process was repeated 3 times until white precipitation ceased. The resultant gel at the bottom of the tube was mixed with 1 mL of anhydrous DMSO. To the dried samples, 200 µL of DMSO was added. Three hundred µL of the DMSO/NaOH base were added, followed by 100 µL of methyl iodide. The sample vial was vortexed and then mixed using a shaker for 15 mins. The reaction was then quenched using 2 mL of LC-MS grade water. Two milliliters of dichloromethane was then added, and the solution was mixed vigorously for 30 secs to extract the permethylated glycans. The sample was then centrifuged at 3000 X g for 1 min to induce phase separation. The upper water layer was removed, and an additional 2 mL of LC-MS grade water was added. This process was repeated 3 times. The lower layer was then transferred to a clean glass vial. The dichloromethane layer was then dried under a stream of nitrogen. The dried sample was then resuspended in 20 µL of MeOH and used for MALDI-TOF/TOF and ESI-MS/MS analysis. MALDI-TOF/TOF analysis was done using a TOF/TOF 5800 MALDI MS (AB Sciex) and ESI-MS/MS analysis was done using an Orbitrap Fusion Tribrid mass spectrometer (Thermo Fisher).

### Sialic acid Linkage Determination

Permethylated O-glycans were dissolved in 50:50 MeOH:H2O with 1 mM lithium carbonate. The samples were injected directly into an Orbitrap Fusion Tribrid mass spectrometer (Thermo Fisher). MS^n^ experiments were then performed to isolate and fragment the penultimate galactose residue (m/z 211.1) adjacent to the sialic acids. Analysis was performed in the ion trap, using quadrupole isolation. Diagnostic fragment ions were then identified to determine linkage as outlined previously by Anthony et al. (Anthony et al., 2008).

### Mass Spectrometry Data Analysis

MS/MS analysis was done using a data-dependent scan method using CID fragmentation. N- and O-glycans were analyzed using GlycoWorkbench 2.0 and Data Explorer software for MALDI data and Freestyle software for ESI-MS/MS data. Glycolipid data was analyzed using Freestyle software, GlycoWorkbench 2.0, and Django software.

### Formation of Oviduct Epithelial Cell Spheroids

Primary oviduct epithelial cells were collected as described above. Collected OECs were washed in PBS for 1 min at 300 x g. Cells were resuspended in 1 mL of TAGP and disaggregated by gently pipetting 10 times with a P1000 pipette. Cells were brought to a 5 mL volume and washed again at 300 x g for 1 min. The OECs were then resuspended in 1 mL of TAGP and then disaggregated gently by passing the cells through a 23-gauge needle ten times. The cells were then brought to approximately a 1 million cells/mL concentration using a hemocytometer and 200 μL were transferred to 60 mm Petri dishes that contain microEB arrays (MicroSurfaces) with circular inserts that are 200 μm in diameter. The OECs were incubated at 39°C in 5% CO2 for 2 hr to allow the OEC spheroids to form. Following incubation, the OEC spheroids were selected based on shape and size and transferred to a wash Petri dish with TAGP before being used in their respective experiments.

### Staining of Oviduct Epithelial Cell Spheroids

The OEC spheroids that were used for staining were incubated with Hoechst 33342 (Invitrogen) before the aggregation step. Once the aggregates were selected and washed, these were fixed in 4% paraformaldehyde for 30 min at room temperature. The OEC spheroids were then washed in dmTAGP and blocked in 5% goat serum with 0.1% Triton X-100 in PBS for 1 hr at room temperature. The OEC spheroids were then incubated with hybridoma supernatants raised against the extracellular regions of porcine Siglecs-1, -3, -5, and -10 overnight at 4°C. The hybridoma supernatants were kindly provided by Dr. Javier Dominguez from the National Institute for Agricultural and Food Research and Technology (INIA) in Madrid, Spain (Álvarez et al., 2015; Escalona et al., 2014, 2015; Revilla et al., 2009). Some aggregates were also stained with α-tubulin (1:200) (Sigma-Aldrich) or E-cadherin (1:100) (Biorbyt) antibodies. Following incubation, the spheroids were washed again in TAGP and incubated with goat anti-mouse Alexa Fluor 488 (1:1000) for Siglec hybridoma supernatants or goat anti-rabbit Alexa 488 (1:1000) and goat anti-mouse Alexa 568 (1:1000) for E-cadherin and alpha-tubulin respectively. Non-specific binding was assessed by incubating the OEC spheroids with IgG isotope control and their respective secondary antibodies. A list of the antibodies used for staining can be found in **Supplemental Table 2**. Some spheroids were also pre-incubated for 30 min at 37°C with 400 μL of sperm stained with Syto Deep Red at a concentration of 1.25 million/sperm per mL before the fixation protocol. All experiments were performed in triplicate in different biological samples. Stained OEC spheroids were mounted in PBS in a glass bottom 96 well plate and imaged using an LSM 700 confocal microscope (Zeiss).

## Results

### Siglec mRNA and proteins are found in cells from the porcine isthmus

To determine the presence of Siglecs in the oviduct at different stages of estrous, the isthmus retrieved from pre-pubertal gilts and cycling post-pubertal multiparous sows were analyzed using end-point RT-PCR. The results indicated that there are 8 Siglecs (Siglec-1, -2, -3, -4, -10, -11, -14, and -15) expressed in the isthmus (Fig. 2A). However, post-pubertal gilts in diestrus showed an absence of Siglec-10. Out of the 8 Siglecs, five possess immune-inhibitory motifs (Siglecs-2, -3, -5, -10, and -11), two possess immune-activating motifs (Siglecs-14 and -15), and one lacks any immunoregulatory motifs (Siglec-1). This is significantly higher than the number of Siglecs reported in other reproductive tissues. For instance, the human endometrium has been reported to express two immune inhibitory Siglecs (Siglec-10 and -11) and one immune activating Siglec (Siglec-16) (Tecle et al., 2019). Through end-point RT-PCR we also found the expression of activating (DAP10 and DAP12) and inhibitory (SHP1 and SHP2) downstream Siglec signaling molecules in the isthmus region of the porcine oviduct regardless of the stage of estrous cycle **(Fig. 2B)**. This implies that the oviduct Siglecs are likely functional and capable of local immune regulation if activated. The high number of Siglecs in the oviduct suggests an intricate regulation of the local immune microenvironment with a bias towards immune suppression.

**Figure 1.**
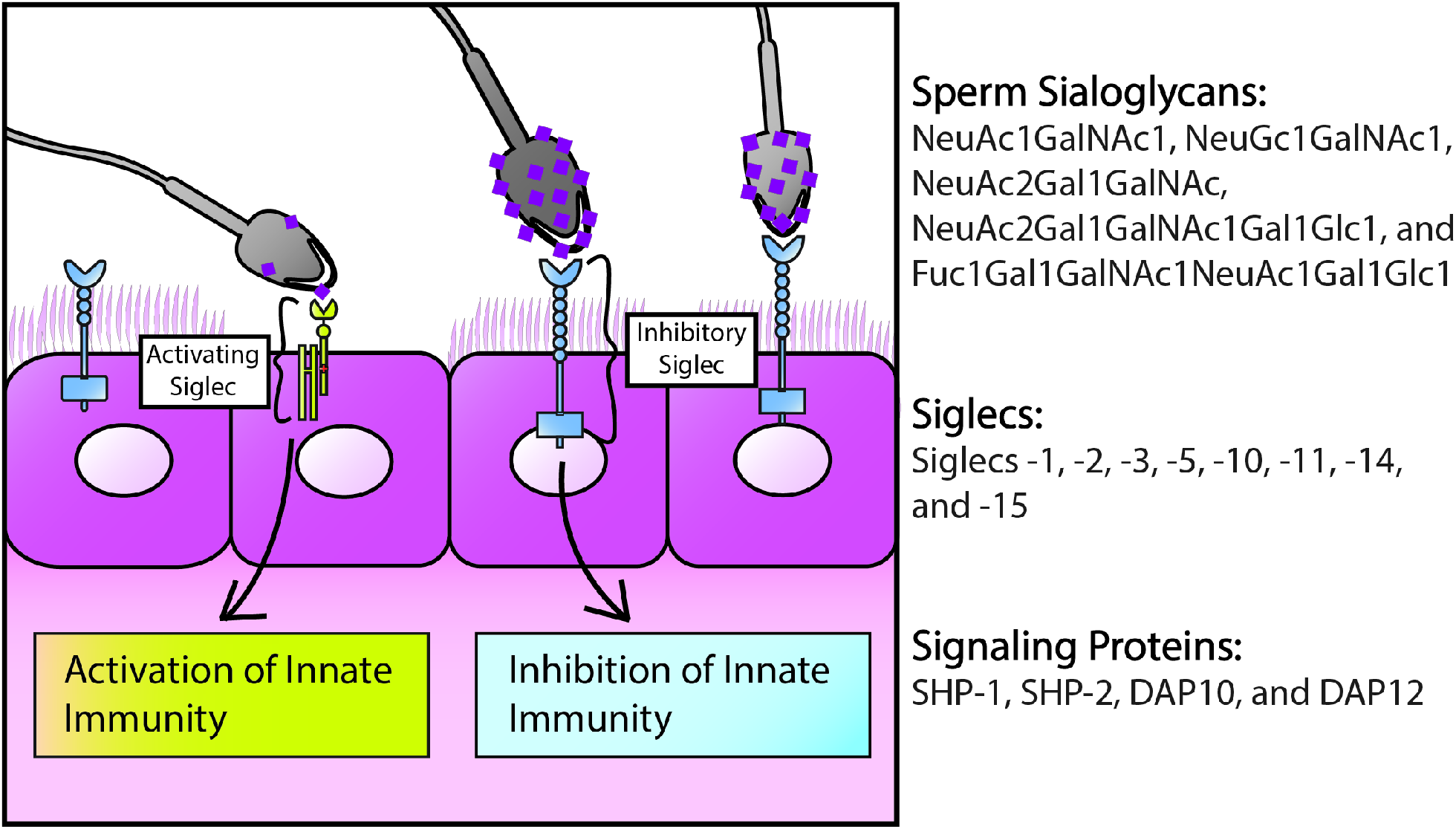
Graphical Abstract. Proposed model of interaction between Siglecs present on oviduct epithelium and sialylated glycans from porcine sperm

**Fig 2.**
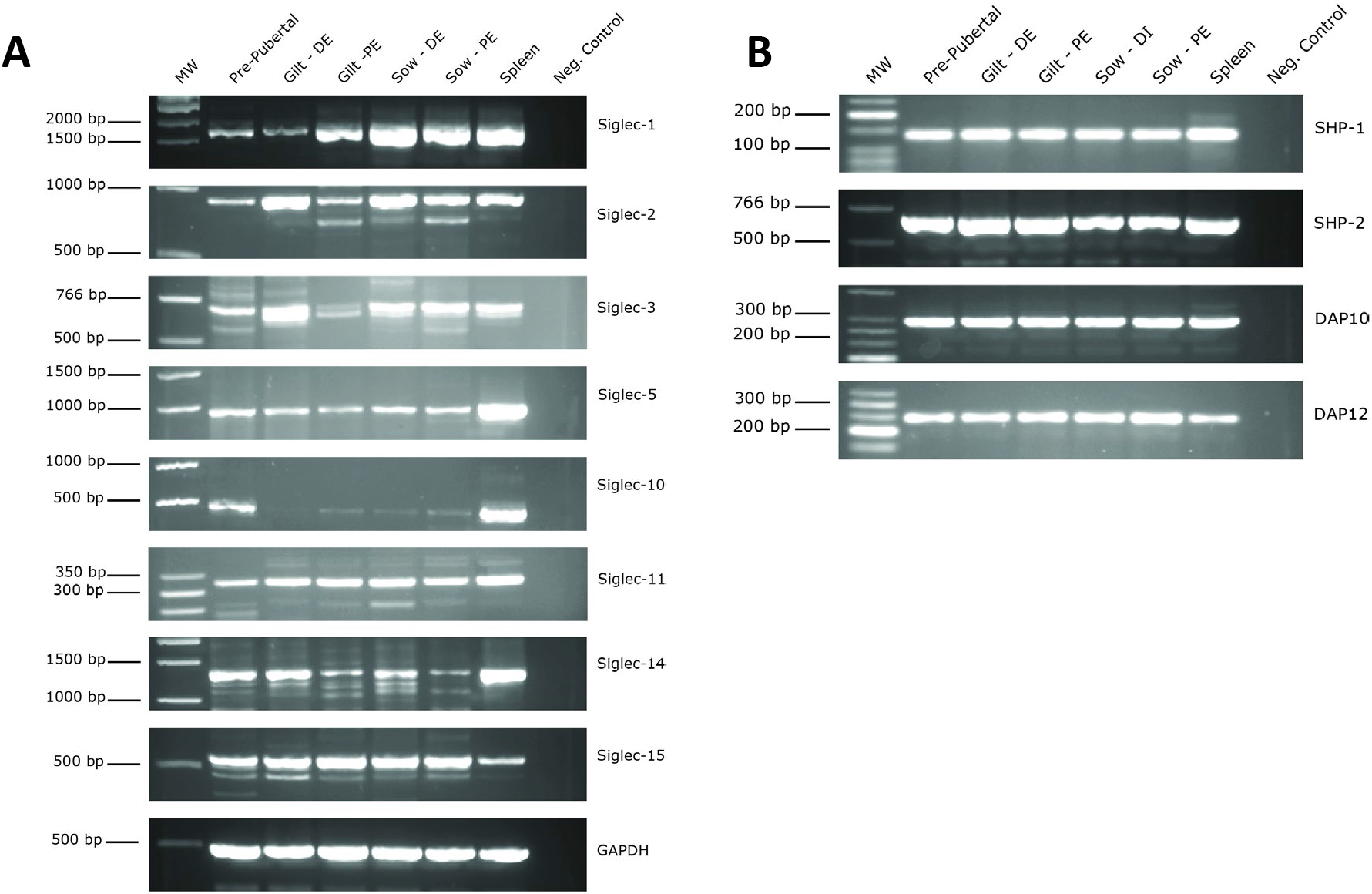
Characterization of Porcine Siglec Gene Expression. Representative results of RT-PCR gels for **A)** Siglecs and **B)** their downstream signaling molecules. Groups include pre-pubertal gilts and diestrus (DE) and proestrus (PE) post-pubertal gilts and sows. The marginal zone of the spleen was used as a positive control. The negative control is a no-template control. GAPDH was used as a loading control for all samples. The first column is either a 1 kilobase or a low molecular weight DNA ladder.

To assess Siglec protein localization, we used oviduct epithelial cells spheroids. Following a 2-hr incubation of primary isthmic oviduct epithelial cells on microEB arrays at 39°C, we were able to efficiently form three-dimensional oviduct epithelial cell spheroids **(Fig. 3)**. The cell aggregates were spherical with an average diameter of 200-250 µm. Immunofluorescent staining demonstrated that the spheroids were primarily formed by E-cadherin positive epithelial cells **(Fig. 3B)** and that ciliated epithelial cells were also present **(Fig. 3A.)** The ciliated epithelial cells were located primarily on the outer region of the spheroids with the cilia facing outward **(Fig. 3B)**. This indicates that the spheroids conserve their apical-basal polarity with the outer cell composition similar to the luminal surface of the oviduct when re-aggregated in vitro. This is important because cilia are known to play a major role in sperm adhesion to the oviduct epithelium (Ardon et al., 2016) **(Supplemental Fig. 1)**. The spheroids are capable of sperm-adhesion interactions as they would normally *in-vivo* **(Fig. 3C)**. Lastly, the oviduct epithelial cell spheroids showed positive signals when stained with Siglec hybridoma supernatants for Siglecs-3, -5, and -10 **(Fig. 4C - E)**. Surprisingly, unlike our PCR results, we were unable to detect Siglec-1 protein in the OEC spheroids **(Fig. 4B)**. This could mean that Siglec-1 mRNA is not translated to a degree that is detectable with immunofluorescence. Alternatively, Siglec-1 protein may be largely in other types of oviduct cells found in the oviduct. This is consistent with another report of Siglecs in the reproductive tract. Siglecs-3, -5, -9, -11, -14, and -16 are present in the cervix (Landig et al., 2019). However, only Siglecs-11 and -16 have been characterized on the cervical epithelium while the rest of the Siglecs are found on different local leukocyte populations (Landig et al., 2019). Lastly, we found that sperm adhesion is co-localized with Siglec-10 staining **(Fig 4F)**, highlighting the probability that Siglec-mediated signaling is induced by sperm adhesion to the oviduct.

**Fig 3.**
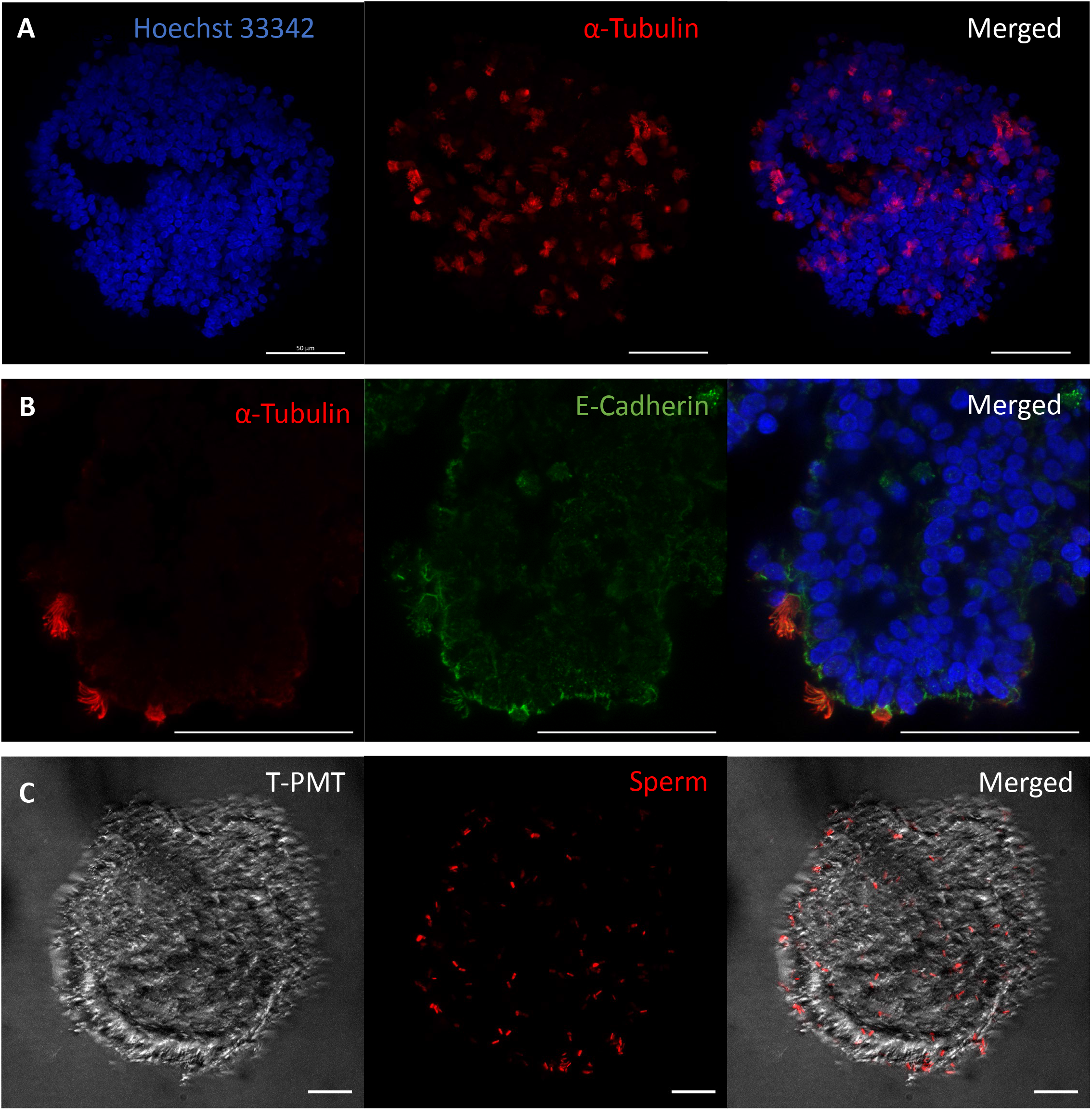
Sperm Bind to Oviduct Epithelial Cell Spheroids: **A**) Oviduct epithelial cell spheroids stained with Hoechst 33342 (blue) and with α-tubulin (red) a marker for cilia. **B)** Cross section of oviduct epithelial cell spheroid stained with Hoechst 33342, α-tubulin, and E-cadherin (green) an epithelial cell marker. **C)** Sperm stained with Syto Deep Red (red) adhering to oviduct epithelial cell spheroid. T-PMT is a photomultiplier for transmitted light. Scale bar is 50 μm.

**Fig 4.**
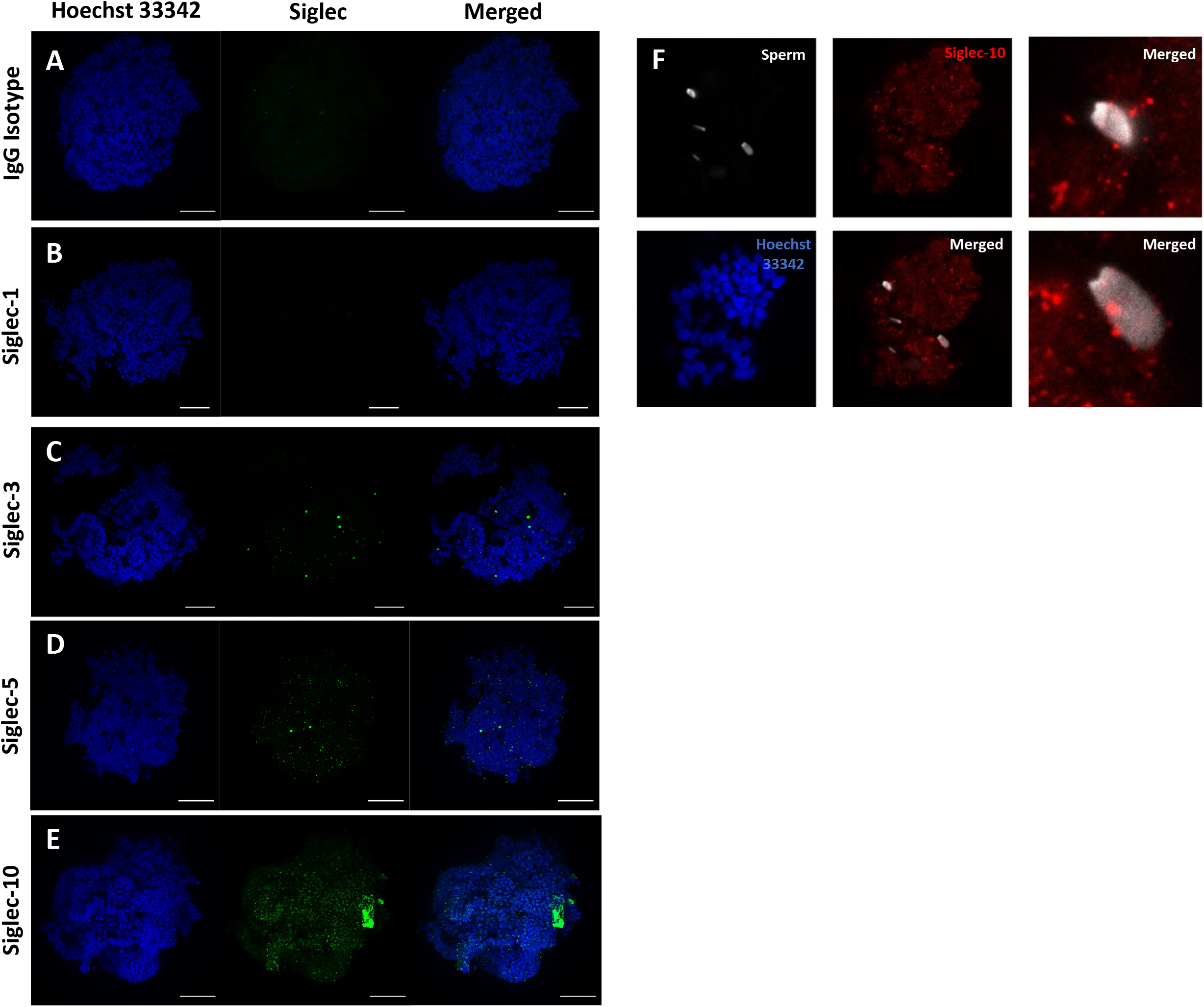
Siglec Localization on Oviduct Epithelial Cell Spheroids. Oviduct epithelial cell aggregate stained with Hoechst 33342 (blue) to stain cell nuclei and with Siglec mouse hybridoma supernatants for **B)** Siglec-1, **C)** Siglec-3, **D)** Siglec-5, and **E)** Siglec-10 were utilized to localize porcine Siglecs on OEC spheroids. **A)** IgG isotype antibody with secondary antibody was utilized to control for non-specific binding. Scale bar is 50 μm. **F)** Sperm binding colocalization to Siglec-10.

### N-glycomics analysis did not identify sialylated structures in porcine sperm

To determine if sialic acid-containing glycoforms were present within the sperm samples, N- and O-glycomics and glycolipidomic analyses were performed. The N-glycans identified include oligomannose, complex and hybrid types (Fig 5A). These included terminal galactose-based sugars as well as terminal acetylhexosamine sugars. N-glycomics analysis by MALDI-MS and ESI-MS/MS did not identify any sialic acid-containing glycans. Lewis structures were identified in low abundance, which has been reported previously on human sperm (Pang et al., 2011). Bisecting GlcNAc, a β1,4-linked GlcNAc attached to the core β-mannose residue and previously noted on human sperm (Pang et al., 2007), was also identified in porcine sperm and confirmed by MS/MS analysis. The most abundant N-glycan was determined to be (Fuc)_1_ + (Man)_3_(GlcNAc)_2_, which makes up approximately 20% of the N-glycome. Excluding (Man)_2_ + (Man)_3_(HexNAc)_2_, which accounts for 10% of the N-glycome, the other 5 most abundant glycoforms are complex structures containing core fucosylation **(Fig. 5A)**. A full list of identified N-glycans and their relative abundances is shown in **Supplemental Table 3**.

**Fig 5.**
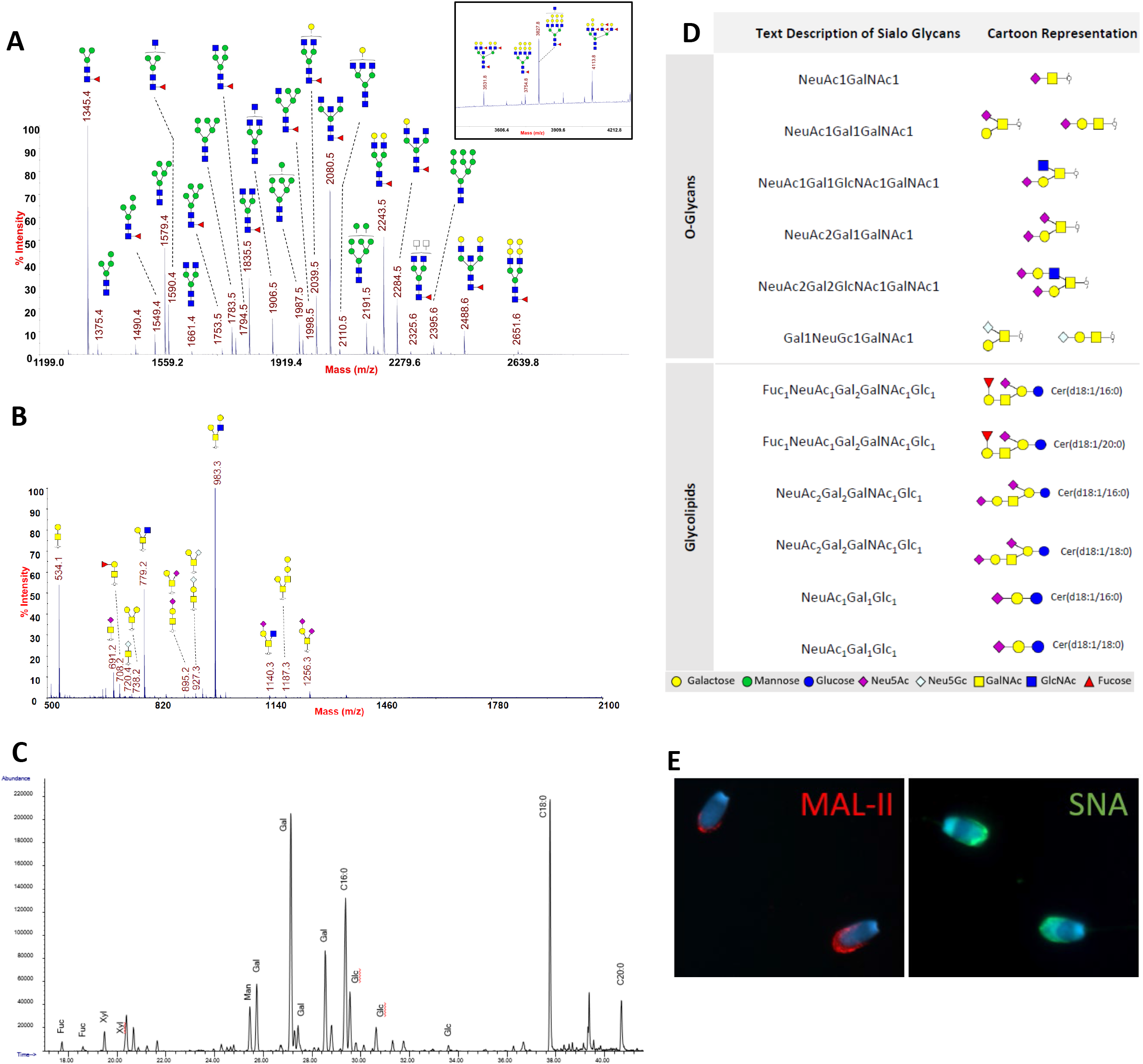
Characterization of Sperm Glycans. **A)** N-glycan MALDI-TOF/TOF results **B)** O-glycan MALDI-TOF/TOF results C) Lipid ESI-CID MS/MS results. Glycosyl composition analysis by GC-MS of TMS-derivatized methyl glycosides **D)** Table showing the sialic acid-terminating glycans profiled on porcine sperm **E)** MAL-II and SNA lectins bind to α- 2,3 and α-2,6 linked sialic acids respectively on the acrosome region of boar sperm.

### O-glycomics analysis identified both non-sialylated and sialylated structures with α2,3-linked sialic acids as the dominant linkage

The O-glycomics analysis detected both sialylated and nonsialylated structures. The most abundant glycoform, (Gal)_2_(GlcNAc)_1_(GalNAc)_1_, made up approximately 38% of the O-glycome. Sialylated structures made up approximately 5% of the O-glycome **(Fig. 5B)**. A full list of identified O-glycans and their relative abundance is shown in **Supplemental Table 3**. Sialic acid linkage was determined by ESI-MS^n^, which fragments the glycan down to the galactose residue linked to sialic acid ^[^29]. For all structures, the linkage was determined to be predominantly α2,3 linked. Indication of α2,6 was also noted for all structures in much lower abundance. An example MS^5^ spectra is shown in **Supplemental Fig. 2A**.

### Glycolipid analysis identified 4 unique glycoforms and 8 glycolipids

Glycosyl composition analysis by GC-MS was performed to determine the presence of monosaccharides and lipid chain lengths. This analysis confirmed the presence of mannose, galactose, glucose, and fucose within the sample. Due to the sample processing, N-acetylhexosamines cannot be distinguished from hexosamines. The fatty acid analysis identified lipid chain lengths of C18:0 and C20:0 **(Fig. 5C)**.

Glycolipid analysis via ESI-MS/MS identified 4 unique glycoforms, and 8 glycolipids. Among the 4 glycoforms identified, 3 contained sialic acid, including sialyllactose (Neu5Ac_1_Gal_1_Glc_1_). The most abundant glycolipids identified contained lactose (Gal_1_Glc_1_) glycoforms and ceramide tails. These made up approximately 66% of the glycolipid abundance, and the remaining 33% corresponds to sialylated glycolipids **(Fig. 5D)**. Glycolipid structures were confirmed by MS/MS **(Supplemental Fig. 2B)**

### Sperm Lectin Staining

The lectins MAL II and SNA, which have a preference for α2,3 and α2,6 linked sialic acids respectively, were used to identify the location of sialoglycans on regions of porcine sperm. The lectin staining patterns varied across sperm within the same sample. However, the staining pattern of α2,3 and α2,6 linked sialic acids in most sperm was localized on the apical region of the sperm head **(Fig. 5E)**.

## Discussion

The glycocalyx of cells is a combination of glycoproteins, proteoglycans, and glycolipids that coat the surface of a cell. Often the termini of glycan structures from the glycocalyx are found to be capped by the monosaccharide sialic acid. However, the function of sialylated glycans found in the sperm glycocalyx has not received much attention. They may be important for sperm function, immune recognition, and survival. For instance, sperm sialic acid levels can influence fertility. A study in chickens showed that sperm with less sialic acid result in lower fertility following insemination (Howarth, 1990). Another study that compared the abundance of sialic acid in the semen from normospermic and oligospermic human patients found that the oligospermic patients had more sialic acid on sperm but less sialic acid in the seminal plasma (Levinsky et al., 1983). However, it appears that sperm sialic acid abundance is not linearly correlated with overall fertility. As seen in a study by Ma and colleagues, the addition or the removal of sialic acids from mouse sperm can result in an increased rate of sperm phagocytosis by uterine macrophages (Ma et al., 2016). Therefore, it seems that an intermediate level of sperm sialylation is needed for proper sperm function and survival inside the female reproductive tract.

Glycan complexity can be achieved through differences in monosaccharide composition, anomeric form, linkage, and capping of the terminal glycan with one or multiple sialic acid residues. Each of these glycan modifications can affect the interactions of sialoglycans with their major cognate receptors of cells, the Siglecs (Büll et al., 2021; O’Reilly & Paulson, 2010). Despite reports highlighting the presence of sialylated glycans on the porcine sperm surface (Jiménez et al., 2003; Wang et al., 2018), none have profiled the full porcine sperm sialome. Accordingly, our objective was to profile the porcine sperm glycome to assess if there were known Siglec ligands present. We characterized the N- and O-glycans and glycolipids in porcine spermatozoa **(Fig. 5A-C)**. Our findings revealed similar results to the profiling of human sperm N-linked glycans. For instance, both studies found high mannose glycans, bisecting glycans, and Lewis structures present among the N-glycans of porcine and human sperm (Pang et al., 2007). However, unlike human sperm no sialic acid containing glycans were found among the N-glycans of porcine sperm (Pang et al., 2007). Nonetheless, we did find unique N-linked glycan peaks that were not observed in human sperm (Pang et al., 2007) and O-linked and galactose-based sugars that contain terminal sialic acids which have previously been found in porcine pancreatic islets (Nanno et al., 2020). To our knowledge, we are the first group to characterize the O-glycans and glycolipids on sperm from any species. Among the glycans and glycolipids we identified, there were several high-preference Siglec ligands (O-glycans: NeuAc_1_GalNAc_1_, NeuGc_1_GalNAc_1_, NeuAc_2_Gal_1_GalNAc_1,_ and Glycolipids: NeuAc_2_Gal_1_GalNAc_1_Gal_1_Glc_1_, Fuc_1_Gal_1_GalNAc_1_NeuAc_1_Gal_1_Glc_1_) **(Fig. 5D)**. Furthermore, we also performed linkage analysis of the sialylated glycans to fully characterize the identity of these sialylated structures **(Supplemental Fig. 2A)**. Our results also showed a mixture of α2,3 and α2,6-linked sialic acids although α2,3-linked glycans were more abundant. It is worth noting that the percentage of total sialic acid-terminating glycans is lower than that seen in human sperm, thus highlighting the differences among species.

Compared to the Siglec expression found in the uterine endometrium from humans and mice, we have shown the porcine oviduct expresses a much greater number of Siglecs, especially those involved in immune suppression. The human uterine endometrium only possesses three Siglecs, two of which are immune inhibitory and one immune activating (Tecle et al., 2019). Tecle and colleagues hypothesized that the uterine Siglecs could play a sperm-protective role following insemination (Tecle et al., 2019). However, we speculate that the immune inhibitory signals produced by Siglec signaling do not have a substantial effect on sperm survival in the uterus due to the high number of immune cells that are present in the uterine milieu (Kaeoket et al., 2001) and the rapid immune inflammatory response that takes place during insemination (Katila, 2012). However, unlike the uterus, interactions between the oviduct epithelium and sperm are prolonged (Birkhead & Møller, 1993; Suarez, 2001) and result in the transcription of several immunoinhibitory mediators that can help boost sperm survival (Mousavi et al., 2021). Therefore, we propose that the promotion of sperm survival by Siglecs is more likely in the oviduct microenvironment than in the uterus.

Although this study does not demonstrate that sperm sialoglycans bind to oviduct Siglecs, several pieces of evidence support a functional role for these co-receptors. For instance, mouse sperm can adhere to select Siglec recombinant proteins in vitro (Ma et al., 2012) suggesting that these interactions are likely to also occur in vivo. Furthermore, it is known that sperm oviduct interactions can result in the expression of various immunosuppressive factors (Mousavi et al., 2021). This shows that sperm interactions can alter the signaling in the oviduct microenvironment. Additionally, it has been shown that sperm sialoglycan abundance can affect the rates of phagocytosis by uterine phagocytes (Ma et al., 2016), highlighting the importance of the sperm sialoglycans in immune suppression. We are not the first to propose that sperm-Siglec signaling could aid in sperm survival inside the female reproductive tract (Tecle et al., 2019) but we are the first to propose this in the oviduct. Based on our findings that sperm have sialoglycans on their surface on the region that binds oviduct cells and that 8 Siglecs, 5 of which are immune inhibitory, are found on cells in the porcine lower oviduct, we propose that sperm sialoglycans mediate sperm-Siglec interactions in the oviduct to stimulate a microenvironment that is immunosuppressed and suitable for prolonged sperm storage during the sperm reservoir.

## Conclusion

Sperm that successfully traverse the uterus and enter the oviduct can bind to the isthmic epithelium and form the sperm reservoir. The function of the sperm reservoir is to serve as a site for the prolonged storage of sperm inside the female reproductive tract until ovulation. Sperm generate a phagocytic response in the uterus but those in the oviduct are tolerated by the immune system, resulting in an enhanced lifespan and an increased time window for fertilization (Holt & Fazeli, 2016). Although there is evidence implicating specific adhesion molecules in the formation of the sperm reservoir (Suarez, 2001), the mechanism that results in an immunosuppressed microenvironment is not well understood.

Multiple studies have shown that the surface overexpression of sialylated glycans is a mechanism utilized by cancer cells to exploit Siglec-dependent pathways (Läubli et al., 2022). The resulting immune suppression through these pathways allows cancer cells to avoid immune rejection. Similarly, certain pathogens such as *Neisseria gonorrhea* (Landig et al., 2019) or *hepatitis B virus* (Tsai et al., 2021) can exploit Siglec-dependent pathways to dampen the host’s immune response to improve the probability of infection. In this study, we discovered the presence of various Siglecs in the oviduct and potential Siglec ligands on porcine sperm. We propose that immune regulation of the oviduct microenvironment during the presence of sperm is regulated through a sialo-Siglec-dependent pathway similar to that utilized by cancer cells and certain pathogens. Because there are other adhesion molecules that sperm and oviduct cells engage to promote sperm retention in the oviduct, Siglecs have a minimal function in adhesion. However, more research is needed to fully characterize the role of sperm sialo-Siglec interactions in the induction of an immunosuppressed oviductal microenvironment in the presence of sperm.

## Author Contributions

LM, DJM, and DM devised the project and designed the experiments. LM, LP, AS, and KD performed the experiments. LM, LP, and DJM interpreted the results. LM and LP drafted the first version of the manuscript and LM, LP, KD, DM, PA, and DJM edited the manuscript.

---

**Supplemental Fig 1.**
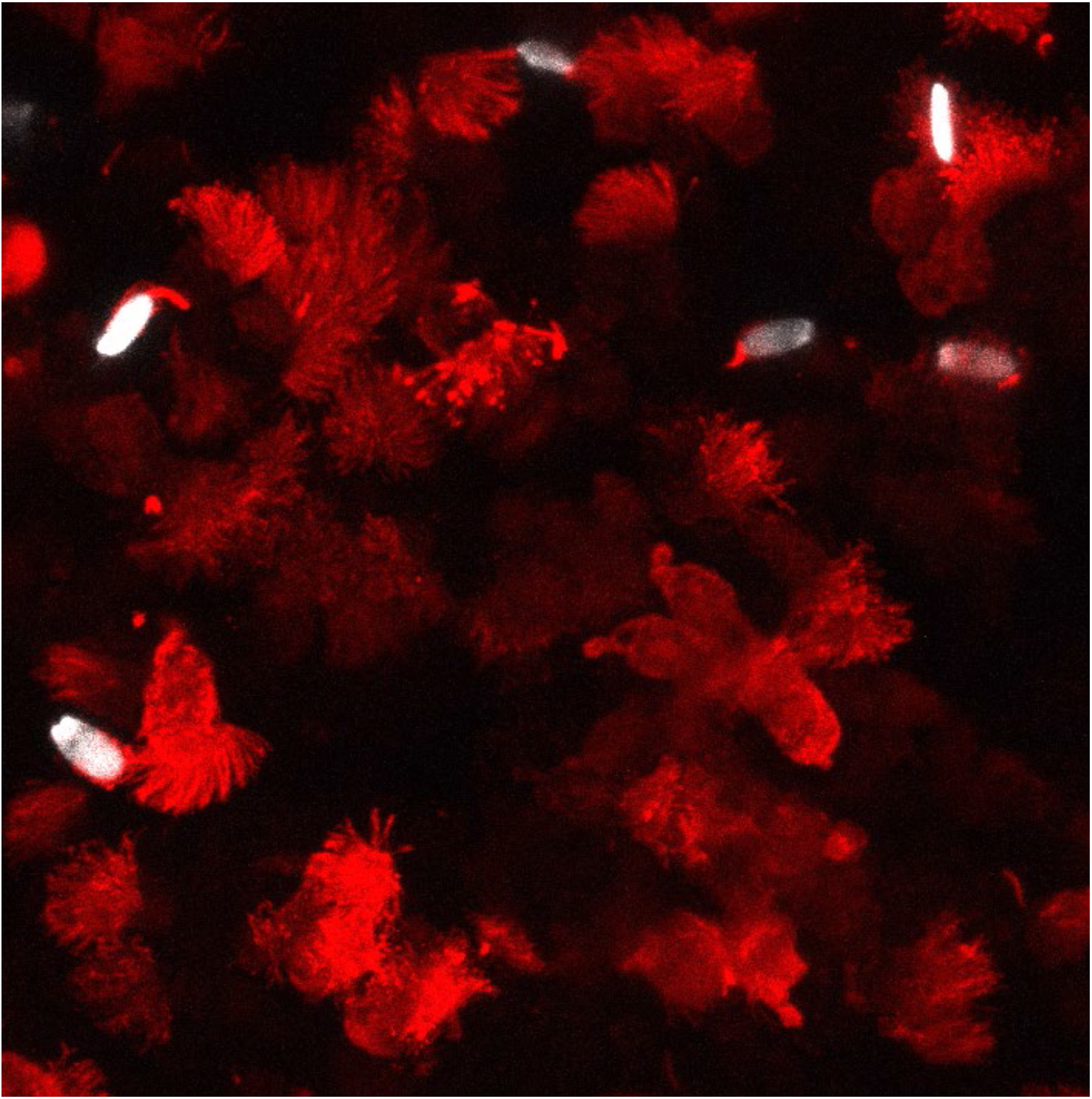
Sperm Binding to Cilia on an OEC Spheroid. Cilia on an oviduct epithelial cell spheroid were stained using an anti-acetyl–alpha tubulin antibody along with Alexa Flour 568 secondary antibody. Sperm nuclei were stained using Syto Deep Red.

**Supplemental Fig 2.**
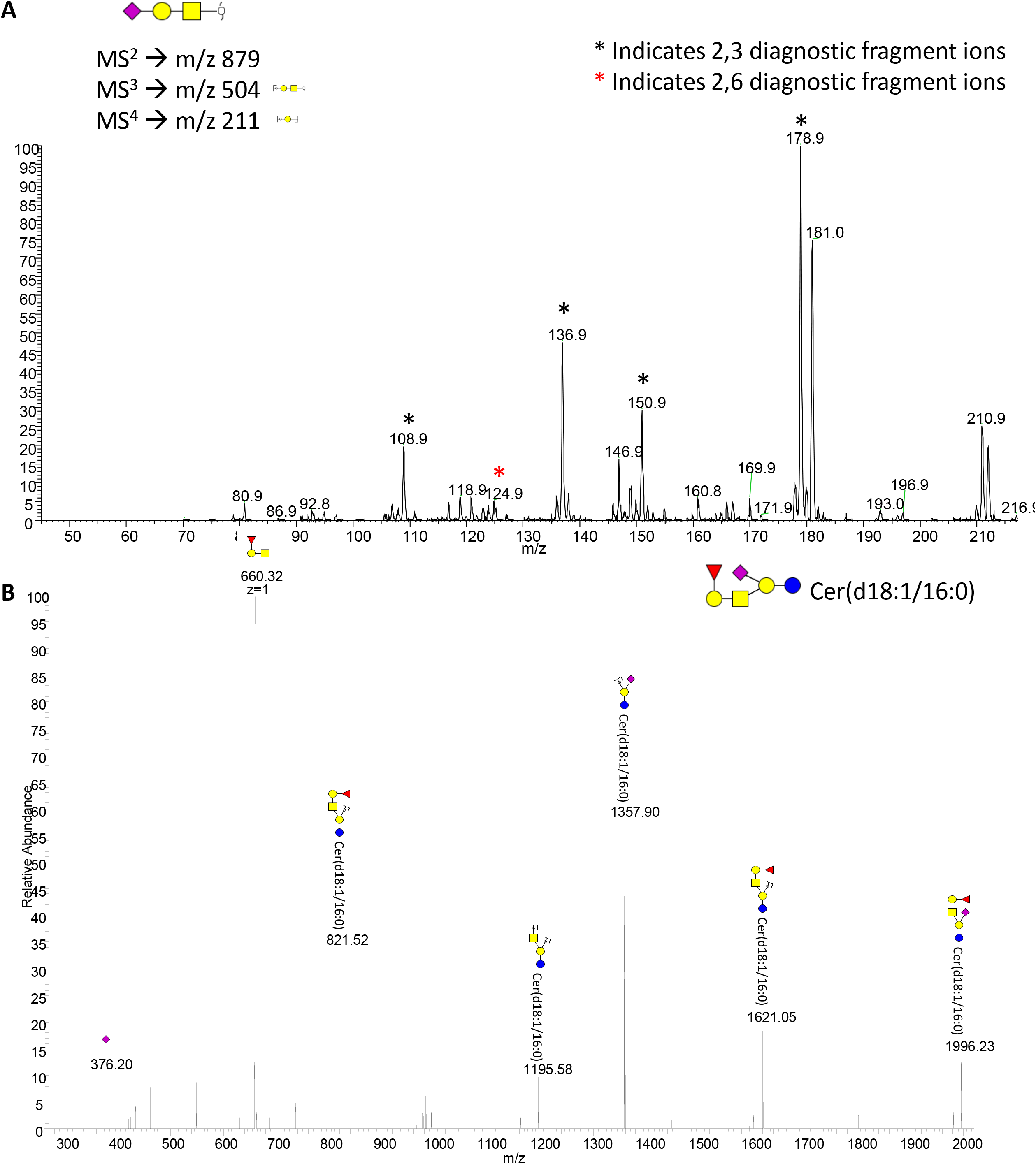
Additional Glycan Spectra. A. Example of MS5 Spectra of Porcine Sperm B. Glycolipid Structures confirmed by MS/MS.

**Supplemental Table 1.**
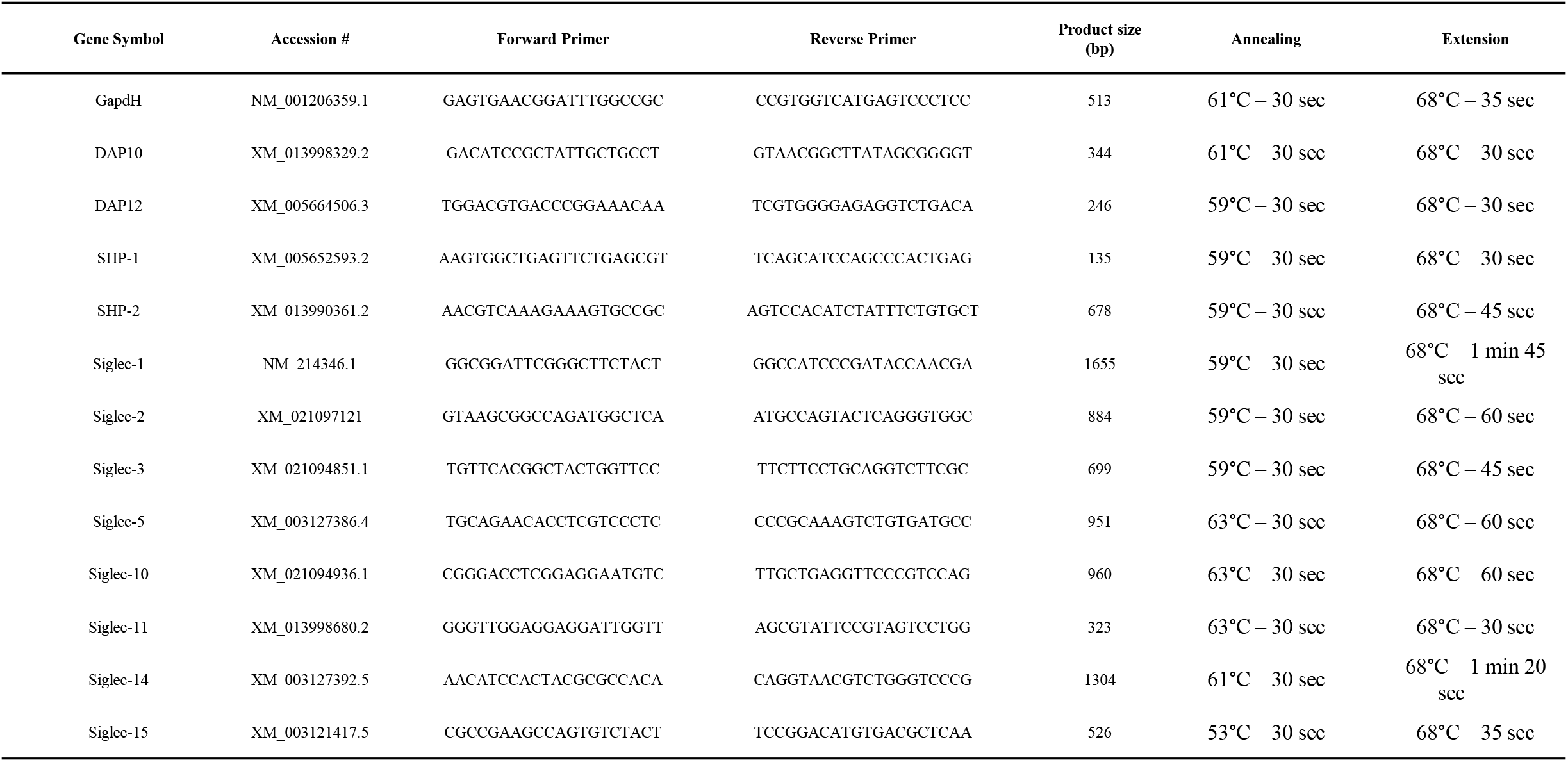
Primer Sequences and End-Point RT-PCR Settings. Gene names and accession numbers were collected from NCBI’s databases. Primer sequences were created using NCBI’s Primer-BLAST tool.

**Supplemental Table 2.** List of antibodies used in immunocytochemistry experiments. * Antibodies were kindly supplied to us by Dr. Javier Dominguez from National Institute for Agricultural and Food Research and Technology in Madrid, Spain.

| Name | Supplier | Cat. # | Host |
| --- | --- | --- | --- |
| CD169 antibody | BioRad | MCA2316GA | Mouse |
| α-Sn 1F1/CR4* | Dr. Dominguez | N/A | Mouse |
| 5D5 Sg-3* | Dr. Dominguez | N/A | Mouse |
| α-Sg5 4F7B7B9* | Dr. Dominguez | N/A | Mouse |
| α-Siglec10 2E9* | Dr. Dominguez | N/A | Mouse |
| Anti-acetyl-α tubulin, clone 6-11B-1 | Sigma-Aldrich | MABT868 | Mouse |
| Alpha Tubulin Polyclonal Antibody | Invitrogen | PA5-16891 | Rabbit |
| E-cadherin antibody | Biorbyt | Orb213706 | Rabbit |
| Mouse IgG Isotype Control | Invitrogen | 08-6599 | Mouse |

**Supplemental Table 3.**
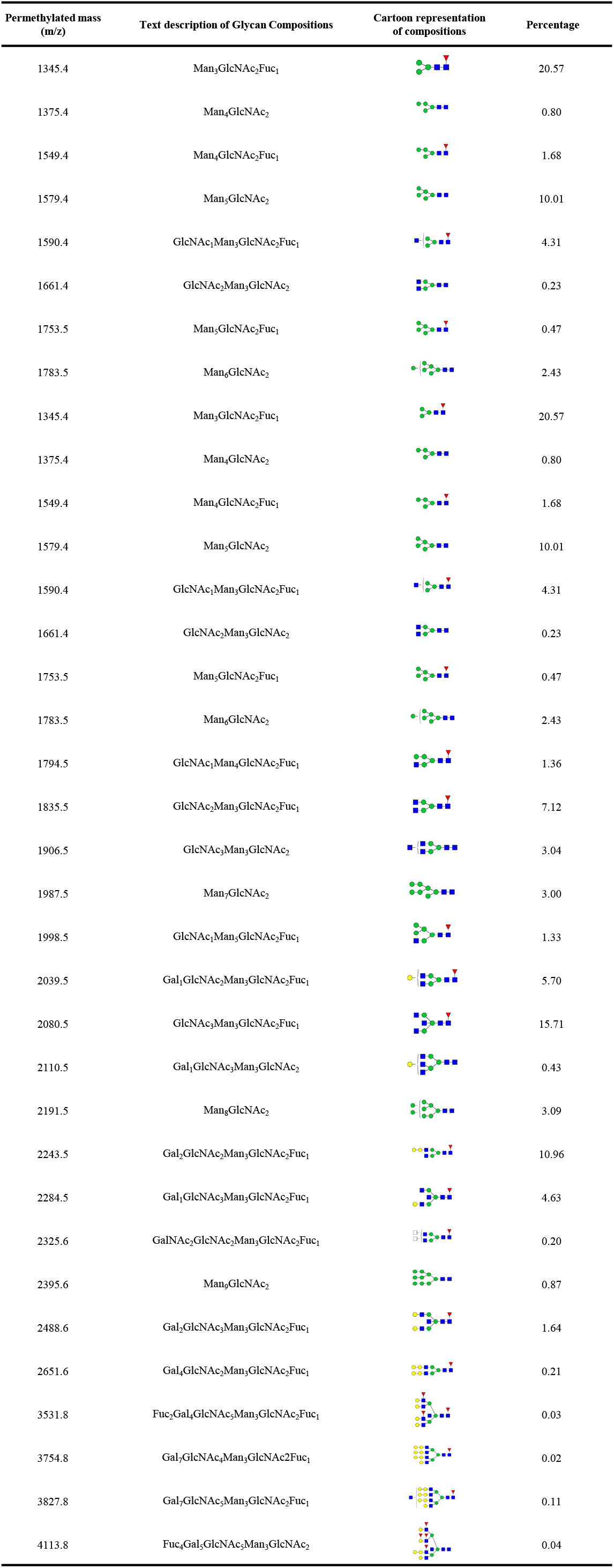
Relative percentage of N-linked glycans of Porcine sperm detected by MALDI-TOF/TOF.

**Supplemental Table 4.** Relative percentage of O-linked glycans of Porcine sperm detected by MALDI-TOF/TOF.

| Permethylated mass (m/z) | Text description of Glycan compositions | Cartoon representation of compositions | Percentage |
| --- | --- | --- | --- |
| 534.1 | Gal<sub>1</sub>GalNAc<sub>1</sub> |  | 25.15 |
| 691.2 | NeuAc<sub>1</sub>GalNAc<sub>1</sub> |  | 4.83 |
| 708.2 | Gal<sub>1</sub>Fuc<sub>1</sub>GalNAc<sub>1</sub> |  | 2.60 |
| 738.2 | Gal<sub>2</sub>GalNAc<sub>1</sub> |  | 0.33 |
| 779.2 | Gal<sub>1</sub>GlcNAc<sub>1</sub>GalNAc<sub>1</sub> |  | 21.65 |
| 895.2 | NeuAc<sub>1</sub>Gal<sub>1</sub>GalNAc<sub>1</sub> |  | 0.61 |
| 927.3 | Gal<sub>1</sub>NeuGc<sub>1</sub>GalNAc<sub>1</sub> |  | 0.78 |
| 983.3 | Gal<sub>2</sub>GlcNAc<sub>1</sub>GalNAc<sub>1</sub> |  | 42.14 |
| 1140.3 | NeuAc<sub>1</sub>Gal<sub>1</sub>GlcNAc<sub>1</sub>GalNAc<sub>1</sub> |  | 0.38 |
| 1187.3 | Gal<sub>3</sub>GalNAc<sub>2</sub> |  | 0.41 |
| 1256.3 | NeuAc<sub>2</sub>Gal<sub>1</sub>GalNAc<sub>1</sub> |  | 1.12 |

